# Testing methods for quantifying Monte Carlo variation for categorical variables in Probabilistic Genotyping

**DOI:** 10.1101/2021.06.25.450000

**Authors:** Jo-Anne Bright, Duncan Taylor, James Curran, John Buckleton

## Abstract

Two methods for applying a lower bound to the variation induced by the Monte Carlo effect are trialled. One of these is implemented in the widely used probabilistic genotyping system, STRmix^™^. Neither approach is giving the desired 99% coverage. In some cases the coverage is much lower than the desired 99%. The discrepancy (i.e. the distance between the *LR* corresponding to the desired coverage and the *LR* observed coverage at 99%) is not large. For example, the discrepancy of 0.23 for approach 1 suggests the lower bounds should be moved downwards by a factor of 1.7 to achieve the desired 99% coverage.

Although less effective than desired these methods provide a layer of conservatism that is additional to the other layers. These other layers are from factors such as the conservatism within the sub-population model, the choice of conservative measures of co-ancestry, the consideration of relatives within the population and the resampling method used for allele probabilities, all of which tend to understate the strength of the findings.

**Highlights:**

1. Two methods for quantifying Monte Carlo variability are tested,
2. Both give less than the desired 99% coverage,
3. The magnitude of possible discrepancy is small,
4. For example an *LR* of 4.3 × 10^11^ could be reported as 1.8 × 10^12^
5. An *LR* of 18 could be reported as 22.

## Introduction

When forensic DNA evidence in presented in court it is valuable, and in many jurisdictions required, that an assessment of the weight of evidence is offered. It is generally accepted that the likelihood ratio (*LR*), or its log, is the best measure to quantify the weight of evidence. The *LR* is usually developed from a number of models and estimates of such things as allele probabilities. Since there are various inputs and decisions along this process there is a range of plausible *LR*s all of which could be judged to be founded on some reasonable set of inputs. The *LR* will be sensitive to changes in modelling assumptions or data to varying degrees depending on the profiles being analysed or compared, the models used, and the assumptions being made. The practise in forensic science is to try and carry out sensitivity analyses that examine the range of the values the *LR* could take given variations in as many inputs of modelling choices and data input as possible [1]. The range of values the *LR* could be assigned (within a reasonable probability interval) will be termed “the plausible range”.

Decision theory utilizes two concepts: probability and utility. Utility describes the expected value of an outcome. For example, the conviction of an innocent person is a catastrophe and it would be universally agreed that the rate of false conviction should be minimized. To round out the set of four outcomes, the conviction of a guilty person is usually accepted to have positive value^5^, the acquittal of an innocent person has positive value, and the acquittal of a guilty person has negative value. To make a decision the decision makers combine probability and utility. Neither the probability nor the utility need to be expressed numerically. However, to optimize your decision making you want the best estimates of both probability and utility. The process is damaged by deliberately biasing either probability or utility.

As a toy example, consider you have a daughter. She has severe eyesight troubles. These troubles are sufficiently bad that they are affecting her access to sport, recreation, potentially affecting her grades, and leading to bullying at school.

There is an operation that could help. This procedure is new, but has been trialled 20 times already. If it works, your daughter’s eyesight will be restored to near to 20/20. However if it fails she may be much worse, even severely visually impaired. The doctors give you the information that the lower bound on the probability of success is 83%. Is this the information you need?

In fact, there has never been a failure out of the 20 previous operations. With only 20 tests it is not possible to say the failure rate is zero. In fact, no number of tests can ever show that the rate of failure is zero. The hospital has commissioned a respected statistician to examine the data and develop advice that can be given to the patients and guardians. This is where the lower bound of 83% comes from. But as one of the primary decision makers for my child I am much more interested in the statistic of 0 failures from 20 operations than in the lower bound. The reason for this is that I need two pieces of information to make this decision: the probability the operation will succeed, and the consequences of both the positive and negative outcomes.

None of the three factors (the probability, the utility of a positive outcome, or the utility of a negative outcome) are known with certainty. We cannot say the probability of the operation failing is exactly 0, nor can we exactly quantify the gain or loss in sight although we have some idea of the range of plausible outcomes. But we do not need absolute certainty in any of these factors to proceed. We all make decisions under such circumstances every day, usually not as serious as the one under discussion, but decisions, nevertheless.

The purpose of introducing this example was to motivate a question that has challenged us for over 20 years. Why do we deliberately bias forensic DNA statistics downwards? This downward bias is termed “conservatism.”

We believe the basis for this is something like: “We consider the conviction of an innocent person a very terrible event and would wish to avoid it whenever possible.” This is a completely rational statement. But it is a statement about utility not probability. What it says is that the utility of a wrongful conviction is strongly negative. A rational response to this equally rational utility statement is to demand a very high standard of proof before convicting anyone [2]. This is only very serendipitously achieved by deliberately biasing the *LR*.

Practice in criminal law courts tends towards an assessment of the lower tail of the plausible range of the *LR*. That represents a deliberate downward bias or in forensic science, conservatism. The desire for conservatism is not necessarily the case for non-criminal matters such as body identification or civil paternity cases, however we do not focus on those cases in our work.

There are a number of inputs into the modelling process used in modern probabilistic genotyping (PG) systems used for the interpretation of forensic DNA profiles. A non-exhaustive list would include:

1. The population genetic model used,
2. The allele probabilities,
3. The value used for the coefficients (i.e. F_ST_) used in the population genetic model.

The three factors listed above would be common to most methods for interpreting forensic DNA profiles and have been previously studied.

The population genetic model used in most PG software follows Balding and Nichols [3] or close variants and is sometimes referred to as NRC II [4] recommendation 4.2. As applied this has been shown to give conservative (high) estimates of the match probability much of the time [5-8].

Few PG take direct account of the uncertainty created when alleles are sampled. Some implement the method of Balding [9] which is not strictly a quantification of sampling uncertainty but which does tend to produce conservative estimates when the population database size is small. STRmix^™^ implements a Bayesian approach called the Highest Posterior Density (HPD) which has been shown to give very good sampling properties [10, 11]. For this system the coverage is close to 99% if *α* = 0.99 is chosen.

For all probabilistic genotyping systems there would be additional factors specific to the modelling of dropout, drop-in, and peak height. Depending on the complexity of the model, it may also include parameters to describe:

- The amount of template DNA each contributor has donated,
- The level of degradation to which each contributor’s DNA has been subjected,
- The efficiency of amplification for each locus examined,
- The amount of peak height variability for alleles,
- The amount of peak height variability for a number of stutter peak types,
- The inter-profile amplification effect for multiple PCR replicates,
- The relative amplification efficiencies of different profiling kits or laboratory processes.

While all these factors can be treated as continuous variables, if they were divided into categorical brackets the number of combinations of parameter values would quickly become astronomical^6^, and too complex to consider exhaustively testing them all. Note that in statistical terms, all these parameters are considered ‘nuisance variables’ because we don’t care what values they take except that we must consider them in order to assign a probability to what we are really interested in evaluating (in this case the probability of the observed peaks given some proposition) (see for example [12]). The choice is then to attempt to simplify the problem with assumptions, or to implement a system of modelling which approximates integrating out the nuisance variables via a stochastic process. An example of this latter solution is Markov chain Monte Carlo (MCMC) and has been implemented in some probabilistic genotyping tools [13, 14]. The important feature of MCMC that makes the complex problem of DNA profile interpretation tractable is that not all possible combinations of parameter values need to be visited in order to get to the set of values that are contributing most to the integrand.

The ultimate value that is being sought is the *LR*, which can be broken down into the two probabilities which need be to be assigned:

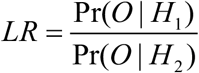

where *H*_1_ and *H*_2_ represent the prosecution and alternate propositions, respectively, and *O* the observed DNA profile. We then consider another nuisance variable (that was not in the earlier list), the genotypes sets, *S*_*j*_, possessed by contributors:

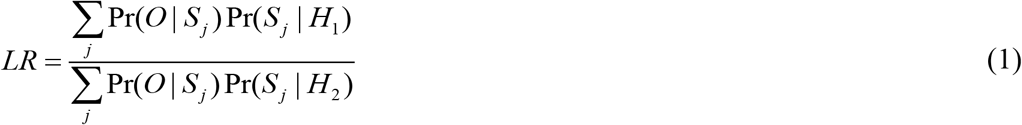

Noting in equation 1 we make the assumption that given the genotype set the probability of the observed data is independent of the proposition. The MCMC provides a sample from the posterior distribution of Pr(*O* | *S* _*j*_), which we term the ‘weight’ for genotype set *j, w*_*j*_. The weights are a categorical variable, i.e. there is a list from which possible genotypes can be chosen, and these have no specific order whereby one leads to another^7^. Within the MCMC, normal treatment of a continuous variable is that a specific value for each variable is trialled within an iteration, and new proposed parameter values are generated by taking small steps away from the currently accepted parameter values. For the categorical parameter of genotype set there are two ways in which they can be treated:

1. Within each iteration the full sum across all genotype sets is carried out
2. Within each iteration only one genotype set is considered, and proposed parameter values include a proposed genotype set, chosen from a list (usually with equal prior probability, although it is also possible to weight the list)

MCMC, being based on a stochastic process, possesses run-to-run variability. The first treatment of genotype sets listed above has the advantage that variability in the weights is naturally taken up into the variability in Pr(*O* | *H*). The disadvantage is that carrying out the MCMC in this manner does not allow easy subsequent *LR* calculation, i.e. if we were interested in the potential DNA contribution of five different people to a complex sample, then the MCMC would need to be run five times, once for each person. There is also a difficultly facilitating human review of the MCMC, as one of the main intuitive diagnostic outputs are the weights [15].

The advantage of the second method of MCMC implementation is that the weights are available to carry out review and subsequent *LR* calculations. The disadvantage is that we may now wish to know how sensitive the *LR* is to the MCMC process of generating weights in this way. In effect the MCMC is now providing values for Pr(*O* | *S* _*j*_) where the integral has been taken across the continuous variables, for each possible value of *S*_*j*_ (one could analogise this to running *J* separate MCMC analyses with a single genotype set fixed in each one). If there is a desire to quantify the lower tail of the *LR* distribution by assessing the variation induced by the MCMC then an understanding of the variance of weights estimated from the MCMC is required. These now have a semi-categorical nature.

Within the MCMC the weights produced are proportional to the probability of the observed data given the genotype sets:

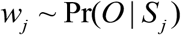

This is sufficient for their use within an *LR* calculation (unless the calculation occurs across more than one number of contributors, in which case marginal likelihoods are required [16]). The values for *w*_*j*_ are obtained by the relative MCMC residence time of genotype set *j*, i.e. the ratio of the number of iterations that genotype set *j* was in the accepted parameter values, divided by the total number of iterations for which the MCMC ran. This is valid as the genotype sets will be accepted and resident in the MCMC for a proportion of time that is relative to the probability of the observed data given that genotype set, compared with others.

If we seek to determine a plausible range for the *LR* given its sensitivity to the MCMC process, then specifically this requires a measure of the variability that may exist in the assigned values for in *w*_*j*_. In this paper we detail two variations on a method that attempts to model variability *w*_*j*_. We test the method by performing coverage tests, i.e. if we consider the sensitivity of the *LR* to only the MCMC assignment of weights, then ideally a 99% lower bound on the *LR* range will be below the average *LR* across many MCMC runs exactly 99% of the time. The results of coverage tests consider only a binary result (i.e. the intervals are either above or below the average *LR*, and the tally of these gives the coverage), and do not give an indication of the level of over- or under-coverage. We also seek to generate a method that provides information on the magnitude of over- or under-coverage when it occurs.

## Method

The weights are assigned from counts of accepted MCMC iterations that contain each genotype set. This may lead to a possible modelling of their variability using a Binomial distribution where successes are accepted iterations and trials are total iterations. There are two issues with using a Binomial method in this manner:

1. The data are correlated, specifically it is well known that adjacent iterations in an MCMC are dependent (and in fact it may be a large number of iterations between independent samples)
2. The data are multi-nominal and have a number of dimensions equal to the number of different genotype sets at a locus (which can be in the 10s of thousands).

We address the first issue by considering how many independent samples occur within the MCMC, a value known as the effective sample size (ESS). Let *x*_1_…*x*_*n*_ be the observations of the likelihood, *p*(*O*|*S*_*j*_) (the lower case *p* signifies a probability density) in a sample obtained from an MCMC run^8^. The mean value of these values is 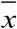. The autocovariance, *γ* (*i*), at *i* is:

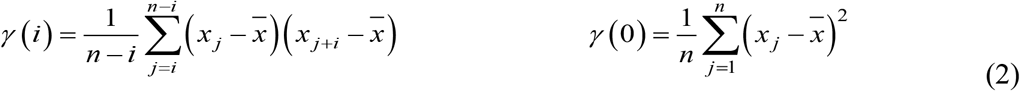

We set *k* as the lowest value of *i* for which *γ* (*i* −1) + *γ* (*i*) ≤ 0 is found.

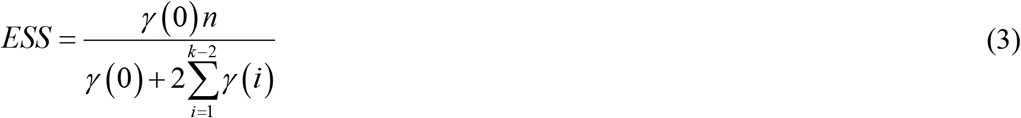

The larger the lag required to reach *k*, the more correlated the data are. This requires a pair of data need to be further apart to be considered independent and consequently the smaller ESS will be. We show two examples of autocovariance values (calculated by equation 2), for an MCMC run that had one chain with a low *k* value and one chain with a high *k* value in Figure 1.

**Figure 1:**
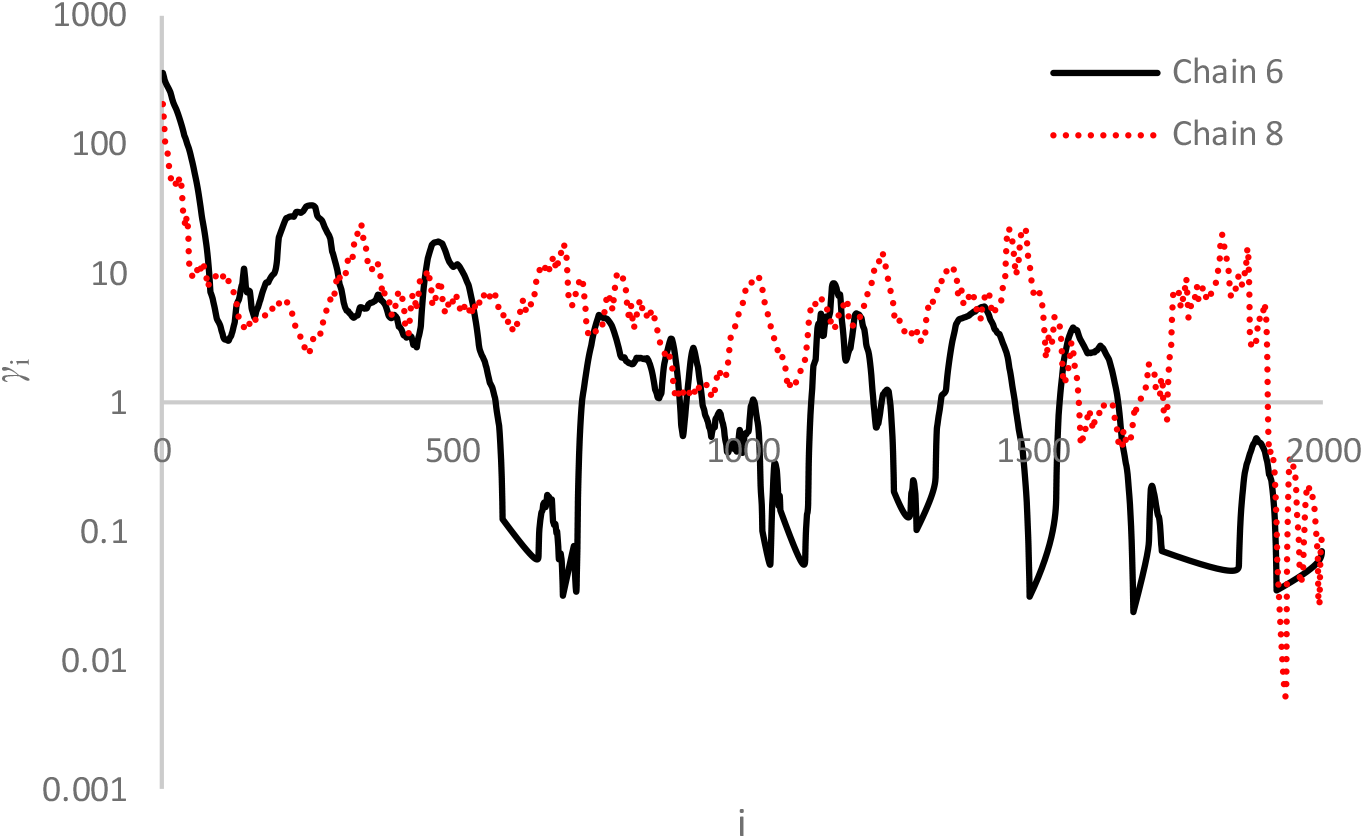
Two plots showing differing autocovariance in two chains from the same deconvolution. Chain 6 (solid black line) has a lower lag value than chain 8 (red dotted line) indicated by chain 6 dropping below 1 on the y axis (axis is shown as log scale) before chain 8.

The approach considers only one chain. In practice, there is usually more than one chain. The methodology we employ, as an approximation to a multichain calculation, is to carry out the ESS calculation on all chains and sum these:

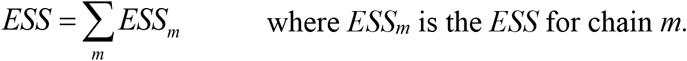

where *ESS*_*m*_ is the *ESS* for chain *m*.

Given that the goal is to find a quantile on a relatively complex function we feel that it is unlikely that an exact theoretical treatment will be available. We propose therefore to assess the coverage empirically.

We set the effective count *EC*_*j*_ for proposed genotype *S*_*j*_ as *EC*_*j*_ = *w*_*j*_ × *ESS*. Consider *EC*_*j*_ to be the number of independent iterations for which genotype set *j* was residence during the MCMC.

In addressing the second issue raised earlier we propose the counts for each genotype to be a multinomial sample with counts. ***EC*** = {*EC*_1_, *EC*_2_,…, *EC*_*N*_} and model these with a Dirichlet distribution.

We compare two variants of our approach.

### Approach 1

In versions of STRmix^™^ from V2.3 ***EC*** was assumed to lead to the joint distribution *Dir*_*N*_ (***EC***_*N*_). Samples (*N* = 1000 by default) are drawn from separate gamma distributions for each of the *J* possible genotype sets *γ* _*j*_ ∼ Г(*EC*_*j*_,1) and resampled weights are set by 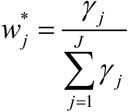 (where the asterisk denotes this is a resampled weight, and so different from the posterior mean value of *w*_*j*_).

### Approach 2

We set a prior distribution for the effective counts *Dir*_*N*_ (**1**_*N*_). The posterior is then *Dir*_*N*_ (***EC*** + **1**_*N*_) and the same Dirichlet-Gamma transform is used for the sampling. The motivation behind this prior is an initial ignorance of any information to use in the prior. However, many other options to model ignorance exist.

In approach 1, the posterior mean of the weights will be same as the weights produced by the MCMC. Using approach 2 the posterior mean is 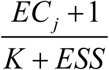, where *K* is the number of different genotype sets with a non-zero weight. This will produce resampled weights with a posterior mean greater than the weight from the deconvolution when 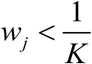, and less than the weight from the deconvolution otherwise.

Using both approach 1 and 2, 11 profiles were deconvoluted *N* = 1000 times each within STRmix^™^ V2.7 [14, 17]. All profiles were GlobalFiler^™^ (Thermo Fisher CA) separated on a 3500 capillary electrophoresis instrument. Seven of the profiles were from the PROVEDIt dataset (two single-source, three two-person mixtures, and two three-person mixtures) and two of the profiles were simulated based on peak height variances observed across the PROVEDIt dataset (one two-person and one three-person mixture) [18]. The remaining two profiles (one single-source and one two-person mixture) were selected because one known contributor gave an *LR* supporting exclusion due to significant dropout across the profile.

For each deconvolution *N* sets of resampled weights were calculated, which were then used to calculate *N LR*s for each profile as per equation 1. *LR*s were assigned using the FBI Caucasian allele frequencies [19] with F_ST_=0.01. Each profile was compared with the known donors (except for one which produced a false exclusion due to an abnormal PCR). This tallied 42,000 deconvolutions for 21 contributor comparisons for each approach. The point estimate (the *LR* calculated using the weights from the MCMC) and lower 0.99 bound on the Monte Carlo variation were recorded using approaches 1 and 2. Note that within the HPD interval only the resampling of weights was carried out, and not any resampling of allele frequencies or F_ST_, so the effectiveness of just the MCMC uncertainty component of the HPD interval could be assessed. The 0.99 lower bound is estimated by drawing a set of *w*_*j*_ values randomly from the gamma distributions described above and calculating the *LR*. This was repeated 1000 times and the lower 0.99 quantile taken as the lower *α* = 0.99 bound.

The average of the *N* point estimates,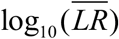, was taken as the best estimate available for the true *LR*. Each of the *N* lower bounds were scored as to whether it was below or above 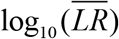. This gives the coverage probability.

## Results

In Figure 2 we show an example of the distribution of 1000 *LR* values obtained for a single analysis of a single, three-person profile.

**Figure 2:**
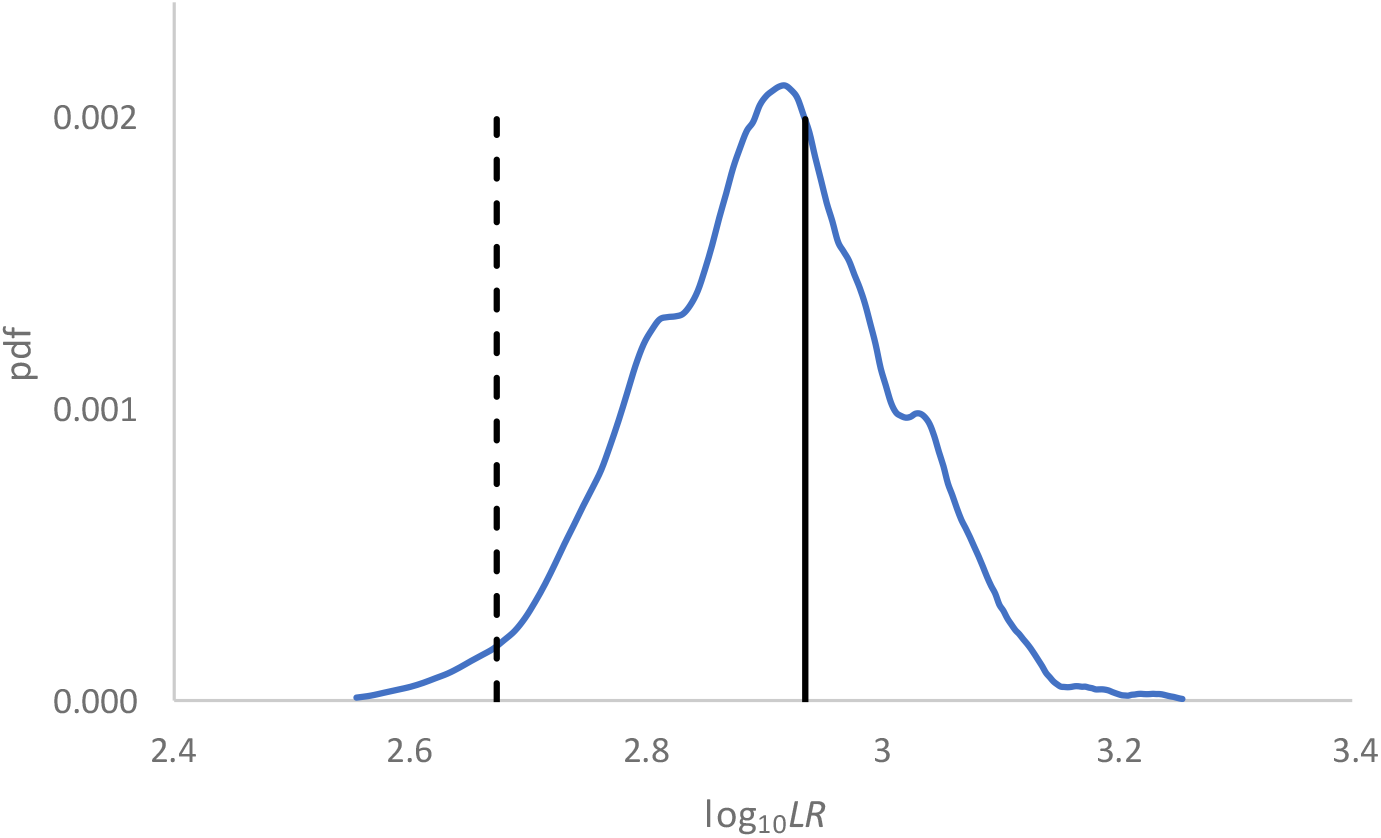
Distribution of log_10_(LR)s for 1000 resamplings of weights for a three-person profile compared with one of the known donors. The dashed vertical line shows the 1-sided, lower bound, 99% interval. The solid vertical line shows the average point estimate.

For each of the *N* runs that each sample was interpreted in STRmix^™^ V2.7 a point estimate and 1-sided, lower bound, 99% interval were generated, as shown in Figure 2 for one known contributor to a three-person profile. We show the distribution of *LR* point estimates and 99% lower bound intervals for the comparisons to the three true donors of a three person mix in Figure 3, i.e. the distributions in Figure 3 are each made up of the 1000 *LR* point estimates (blue distribution) and 1000 99% HPD intervals (red distribution). The solid lines in Figure 3 represent the 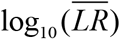, and the dashed line represents the upper 99% of the lower-bound 99% HPD intervals. If performing exactly as desired the dashed line and solid lines in Figure 3 would fall on the same point.

**Figure 3:**
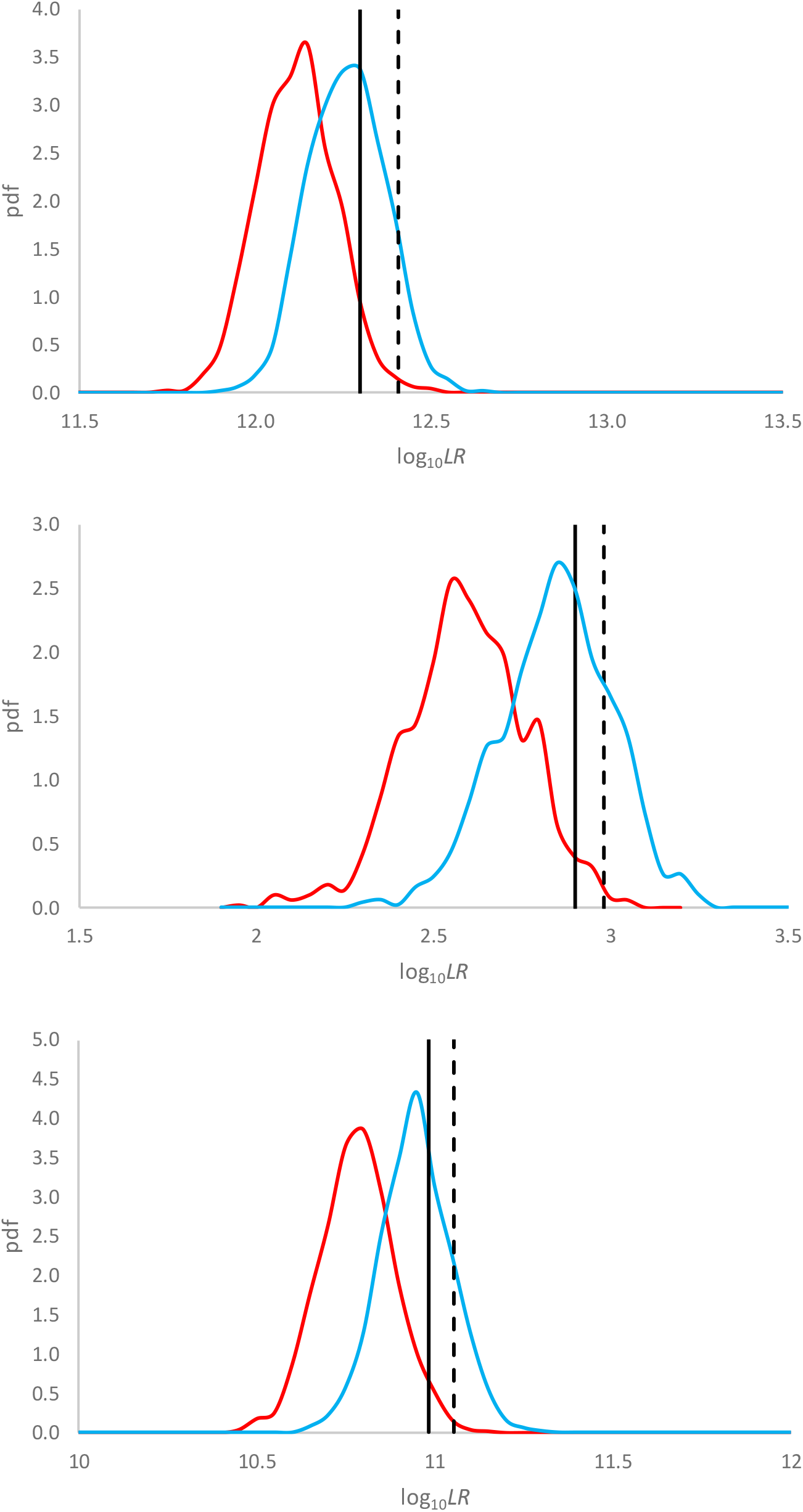
Distribution of point estimate (blue) and 1-sided, lower bound, 99% interval (red) LRs for sample C10 vs the three known donors using approach 1. The solid vertical line is drawn at 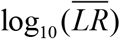. The dashed vertical line shows the 99% upper bound value of the lower bound intervals. If performing exactly as desired the dashed and solid vertical lines would fall on the same point. If these lines coincide then exactly 99% of the of the lower bounds are above the average LR.

Our data show a variety of performances with regards to coverage. Some are below the desired 99%. This should be expected as we are applying it in a complex situation beyond that for which it was derived. Figure 4 and Table 1 give the coverage results for all samples tested using approach 1 and 2 plotted against 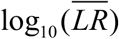.

**Table 1:** 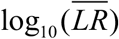 and coverage results for approaches 1 and 2. The discrepancy columns give a measure of how much higher or lower (on a log_10_ scale) the lower bounds would need to be to obtain exactly 99% coverage. The positive values for discrepancy are those where the lower bounds need to be even lower.

| Sample | Contributor | Approach 1 |  |  | Approach 2 |  |  |
| --- | --- | --- | --- | --- | --- | --- | --- |
| | | $\log_{10}(\overline{LR})$ | Coverage | Discrepancy | $\log_{10}(\overline{LR})$ | Coverage | Discrepancy |
| A02 2p | 1 | 6.5 | 93.8% | 0.09 | 6.5 | 94.1% | 0.13 |
|  | 2 | 5.8 | 98.1% | 0.02 | 5.8 | 97.0% | 0.04 |
| A05 2p | 1 | n/a |  |  |  |  |  |
|  | 2 | 30.9 | 81.7% | 0.03 | 30.9 | 89.0% | 0.06 |
| A08 3p | 1 | 8.0 | 88.8% | 0.14 | 8.0 | 93.9% | 0.11 |
|  | 2 | 7.5 | 90.7% | 0.14 | 7.5 | 86.4% | 0.17 |
|  | 3 | 30.2 | 76.1% | 0.11 | 30.2 | 95.9% | 0.05 |
| C06 A 1p | 1 | 22.8 | 100.0% | -0.09 | 22.8 | 100.0% | -0.08 |
| C06 B 1p | 1 | 13.4 | 100.0% | -0.11 | 13.4 | 100.0% | -0.11 |
| C10 3p | 1 | 2.9 | 95.7% | 0.08 | 2.9 | 96.5% | 0.07 |
|  | 2 | 11.0 | 95.1% | 0.07 | 11.0 | 99.0% | -0.01 |
|  | 3 | 12.3 | 92.3% | 0.11 | 12.3 | 96.5% | 0.07 |
| E05 2p | 1 | 17.9 | 91.7% | 0.05 | 17.9 | 92.4% | 0.05 |
|  | 2 | 31.8 | 84.7% | 0.00 | 31.8 | 93.5% | 0.01 |
| sim 3p | 1 | 11.6 | 94.3% | 0.23 | 11.6 | 40.4% | 0.63 |
|  | 2 | 8.5 | 92.5% | 0.11 | 8.5 | 96.4% | 0.06 |
|  | 3 | 1.3 | 96.4% | 0.10 | 1.3 | 99.0% | -0.02 |
| sim 2p | 1 | 16.0 | 100.0% | -0.30 | 16.0 | 99.4% | -0.07 |
|  | 2* | 1.6 | 98.6% | 0.01 | 1.6 | 97.7% | 0.03 |
|  | 3 | 4.1 | 97.8% | 0.02 | 4.1 | 98.0% | 0.02 |
| 1p excl | 1 | -10.0 | 100.0% | -2.00 | -10.0 | 100.0% | -0.46 |
| 2p excl | 1 | -5.8 | 93.1% | 0.30 | -5.8 | 92.1% | 0.26 |

\* Note that contributor 2 to the simulated two-person mixture was a revision of the true contributor's profile who we term contributor 3. This was done in order to lower the LR closer to 1

**Figure 4:**
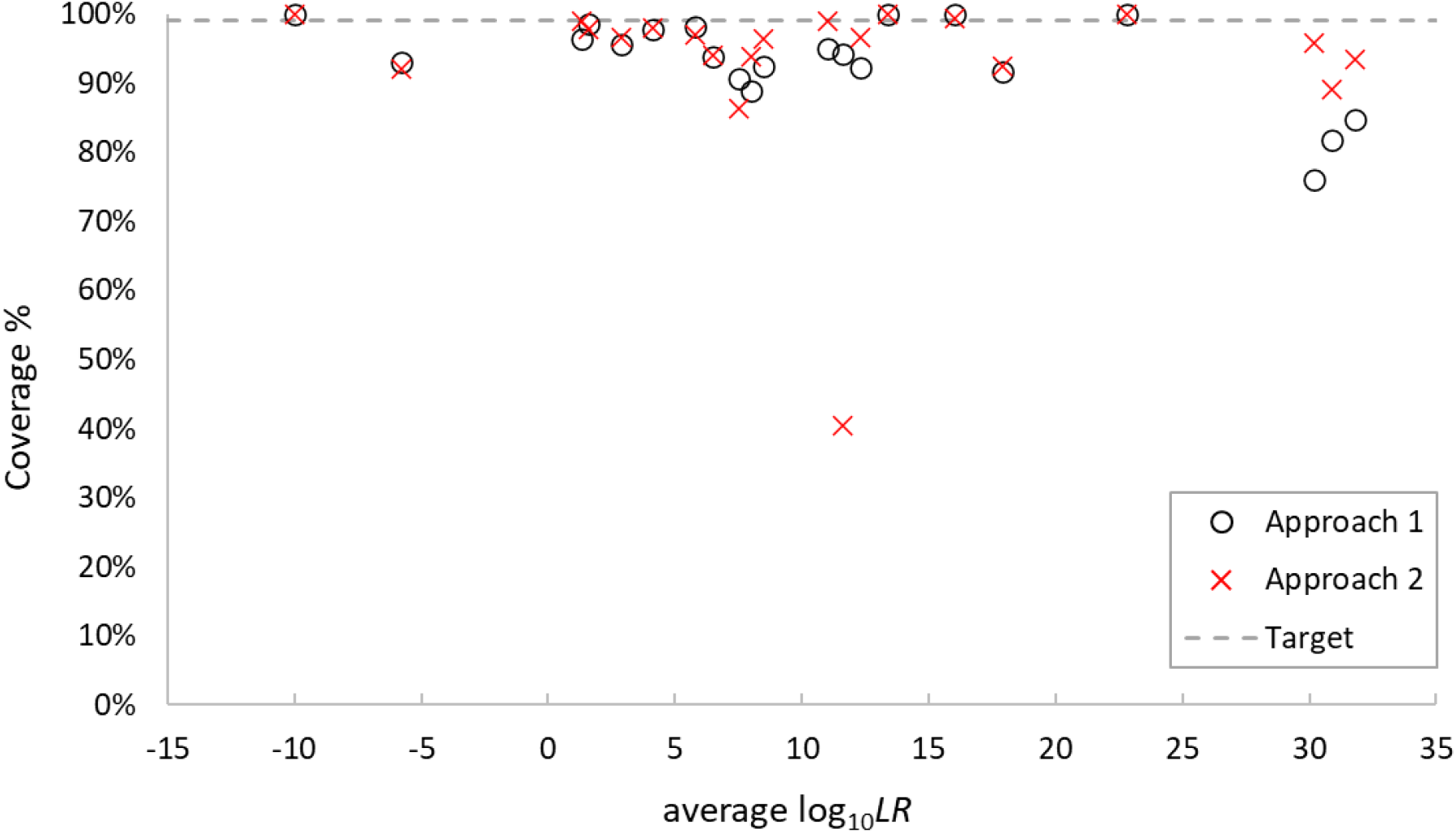
Coverage results for all 21 contributor comparisons plotted against 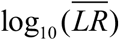. The dashed line is at 99% coverage.

The average coverage using approach 1 was 94% and using approach 2 was 93%. From Table 1 it can be seen that the worst performing individual sample coverage occurred at values of 76.1% and 40.4% using approaches 1 and 2 respectively. These two worst performing coverages were for different samples.

It may be asked how a coverage could be less than 50%. This can only occur for approach 2. This has occurred due to a number of lower bound intervals being above the point estimate for this sample when using approach 2. Recall that the posterior mean of the resampled weights will be greater than the weight from the deconvolution when 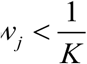. In the sample that displayed the coverage of 40.4% there were three loci where all of the genotype sets that possessed the genotype of the POI in the target contributor position (i.e. those considered in *H*_*p*_) had weight below 1/*K*. The proportion of genotype sets considered under *H*_*p*_ with 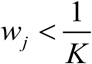 is not the only factor that influences whether the HPD interval will be above the *LR* point estimate, but it is one of the main contributors. Other contributing factors include the absolute number of genotype sets considered under *H*_*p*_ or *H*_*d*_ that also possess 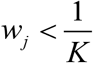

We investigated a number of additional factors (average log(likelihood), Gelman-Rubin convergence diagnostic, and the number of contributors assigned during interpretation) to examine whether any of these had an effect on coverage. We did not detect any relationship between any of these factors and coverage (data not shown).

## Discussion

Examination of Figure 4 and Table 1 suggests that neither approach 1 nor 2 are giving the desired 99% coverage. In some cases the coverage is much lower than the desired 99%. The discrepancy (i.e. the distance between the *LR* corresponding to the desired coverage and the *LR* observed coverage at 99%) is not large. For example, the discrepancy of 0.23 for approach 1 suggests that the lower bounds should be moved downwards by a factor of 1.7 (the inverse log_10_ of 0.23) to achieve the desired 99% coverage.

There are various factors that may account for differing levels of coverage. The more complex profiles will have a greater sample space which can reasonably describe the observed data. If the chains are traversing this large space such that their individual variance is close to the steady state variance (i.e. they are considered converged according to diagnostics) it may still be the case that genotype set residence times are more variable than the ESS method used will cover. This was not seen in our analyses, but may be revealed in larger, and more controlled experiments. One possible solution would be to run the MCMC for an extended number of iterations to produce more stable weights between MCMC runs. However this would also increase the ESS, resulting in tighter intervals. While this is the expected and desired behaviour of the interval, we are unsure of the balance of these two effects with respect to coverage.

An example of these opposite effects is shown in Figure 5, which displays the results in the format of those seen in Figure 3, for sample A08 compared to contributor 3 (the worst of the performing coverages using approach 1). Specifically, Figure 5 shows the distributions of log_10_ (*LR*) point estimates and intervals for a set of standard MCMC analyses, and then those with 10 times the number of iterations. The effects of a reduced variability in the log_10_ (*LR*) can be seen, along with the corresponding reduction in the size of the HPD interval, leading to very similar coverage results regardless of the iterations (76.1% under standard conditions and 74.8% under extended conditions).

**Figure 5:**
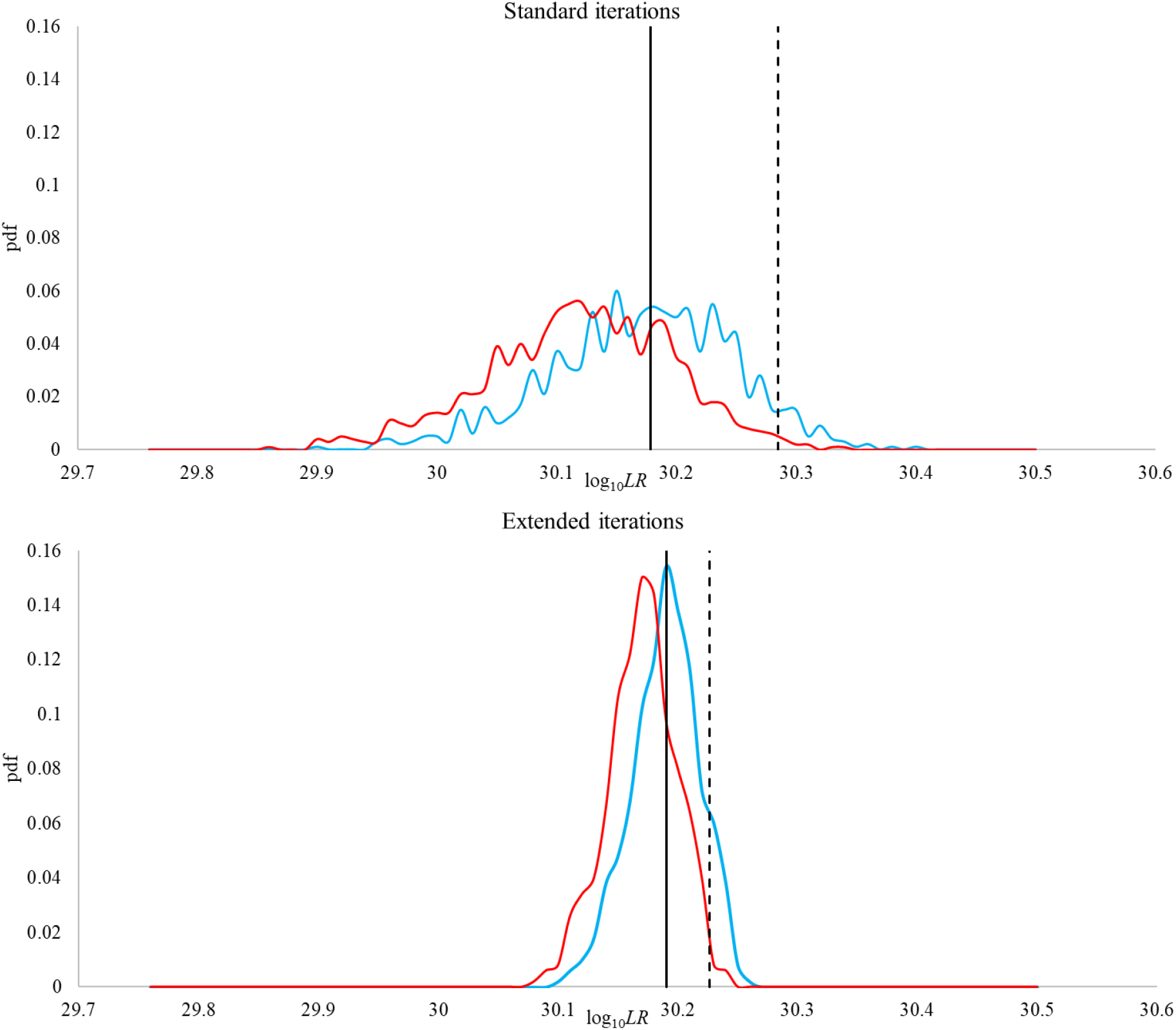
Distribution of point estimate (blue) and 1-sided, lower bound, 99% interval (red) LRs for sample A08 vs contributor 3 using approach 1. The solid vertical line is drawn at 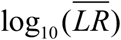. The dashed vertical line shows the 99% upper bound value of the HPD intervals. The top graph is the MCMC run under standard iterations and the lower graph is the same analysis but extending the iterations by a factor of 10.

In our analyses we saw that the lowest coverage was provided for those samples that yielded the highest *LR*, i.e. those that are most resolved and with weights closest to 1. In this situation we would expect that MCMC variability in the *LR* would be low, and that the corresponding HPD interval would be close to the point estimate (which was observed to be true in both instances). Our analysis shows that the method used to determine the interval underestimates the level of MCMC variability in these instances, however the magnitude of the underestimate is slight (as seen in Table 1) and arguably at this level has very little effect on interpretation.

The performance of the method for quantifying Monte Carlo falls short of providing the 99% coverage desired. Although less than desired it does provide a layer of conservatism that is additional to the other layers described in the introduction.

## Acknowledgements

This work was supported in part by grant NIJ 2017-DN-BX-0136 from the US National Institute of Justice. Points of view in this document are those of the authors and do not necessarily represent the official position or policies of their organizations.

## Footnotes

5 We acknowledge valid sociological discussion about how best to treat this situation.

6 Astronomical is an apt description of the size of problem. Even if we considered a relatively coarse breakdown whereby there were 100 different brackets of DNA amount for each contributor, and 100 brackets of possible degradation, and 100 brackets of locus amplification for each locus, and 100 brackets for peak height variabilities, and 100 brackets for replicate amplification, then for a 4-person, 2-PCR GlobalFiler^™^ profile there are 10^66^ combinations of parameter values. This is without even without considering different genotype combinations, which could easily push the number of parameter value combinations over a googol (10^100^). For reference, the number of atoms in the observable universe is only estimated at 10^80^.

7 We are talking about an order in a statistical sense, and not in the sense of a numerical ordering convention that make human reading of the list more acceptable.

8 This process can occur on all iterations, or a thinned selection. Any amount of thinning should in theory provide a similar result, although thinning will tend to lose some power and so can underestimate the ESS. We do not show the mathematics of accounting for thinning in this paper. STRmix^™^ uses an automatically thinned set of values so that the total number of iterations per chain is 100,000.

